# pH-dependent antibacterial activity of N-acetylcysteine against *Staphylococcus aureus* and metabolic alterations induced at neutral pH

**DOI:** 10.64898/2026.06.05.730338

**Authors:** Theresa Schiemer, Claudia Siverino, Vimalnath Nambiar, Kristaps Klavins

**Affiliations:** Institute of Biomaterials and Bioengineering, Faculty of Natural Sciences and Technology, Riga Technical University, Riga, Latvia; Baltic Biomaterials Centre of Excellence, Headquarters at Riga Technical University, Riga, Latvia; AO Research Institute Davos, Davos, Switzerland; Australian National Phenome Centre, Health Futures Institute, Murdoch University, Murdoch, Australia; Centre for Computational and Systems Medicine, Health Futures Institute, Murdoch University, Murdoch, Australia

**Keywords:** N-acetylcysteine, Staphylococcus aureus, pH-dependent antibacterial activity, metabolomics, arginine metabolism

## Abstract

N-acetylcysteine (NAC) is a mucolytic and antioxidant increasingly investigated as an antibacterial agent and antibiotic adjuvant, particularly against biofilm-associated infections. Despite being a weak organic acid, solution pH is rarely reported in the literature, and reported minimum inhibitory concentrations (MIC) for *S. aureus* are highly inconsistent across studies, varying by more than 50-fold. We systematically assessed the pH-dependence of NAC antibacterial activity and investigated how pH-neutral NAC perturbs bacterial metabolism using LC-MS based targeted metabolomics.

Without pH-adjustment, NAC at 200 mM (pH 2.8) acted bactericidal against the *S. aureus* strains USA300 and Mu12, while pH-adjusted NAC solutions had no effect on bacterial growth, even at concentrations approaching its solubility limit. Based on LC-MS measurement, pH adjustment did not measurably degrade NAC but significantly increased dimerization ratios, suggesting that altered ionization state rather than degradation underlies the loss of antibacterial activity at neutral pH. Despite the absence of growth inhibition, pH-neutral NAC induced concentration-dependent metabolic alterations under both planktonic and biofilm conditions. Arginine, cysteine, glycolysis, and TCA cycle were the most affected pathways, with intracellular accumulation of arginine and cystine, together with increased lactic acid production.

Our findings demonstrate that the antibacterial activity of NAC against *S. aureus* is driven by acidification, helping reconcile contradictory MIC reports in the literature. Additionally, pH-neutral NAC alters bacterial metabolism without impairing growth, highlighting its potential to modulate bacterial physiology independently of direct antibacterial activity.

## Introduction

Biofilm-associated implant and fracture-related infections remain difficult to treat because of bacterial persistence, antibiotic tolerance, and infection recurrence (1–3). Staphylococcus aureus, the leading causative pathogen, is therefore a major target for alternative antimicrobial and antibiofilm strategies. (4) Among these, N-acetylcysteine (NAC), a clinically used mucolytic and antioxidant, has attracted increasing interest because of its reported antibacterial, antibiofilm, and antibiotic adjuvant properties (5–7).

Reports on NAC’s effect on the most common cause, *S. aureus*, include minimum inhibitory concentration (MIC), biofilm disruption, and antibiotic synergy. Reported results however show great variability: MICs for *S. aureus* ranged from 2.5 mg/mL (8), to as high as 128 mg/mL (9) for clinical isolates. Biofilm disruption is a common finding in the literature (10), however biofilm-enhancing effects have been found as well (11). For antibiotic combination, synergistic effects (12), and antagonistic effects (13) were reported. Landini *et al.* suggested that these discrepancies might be due to NAC’s often overlooked chemistry, specifically its pH (14). Consistent with this observation, revision of publications between 2016 and 2025 identified that only a limited proportion of studies report solution pH in the title or abstract: fewer than 1% overall and only 3% of studies involving bacteria (Fig. 1a). This lack of reporting warrants closer examination of NAC chemistry and its proposed antibacterial mechanisms.

**Fig. 1.**
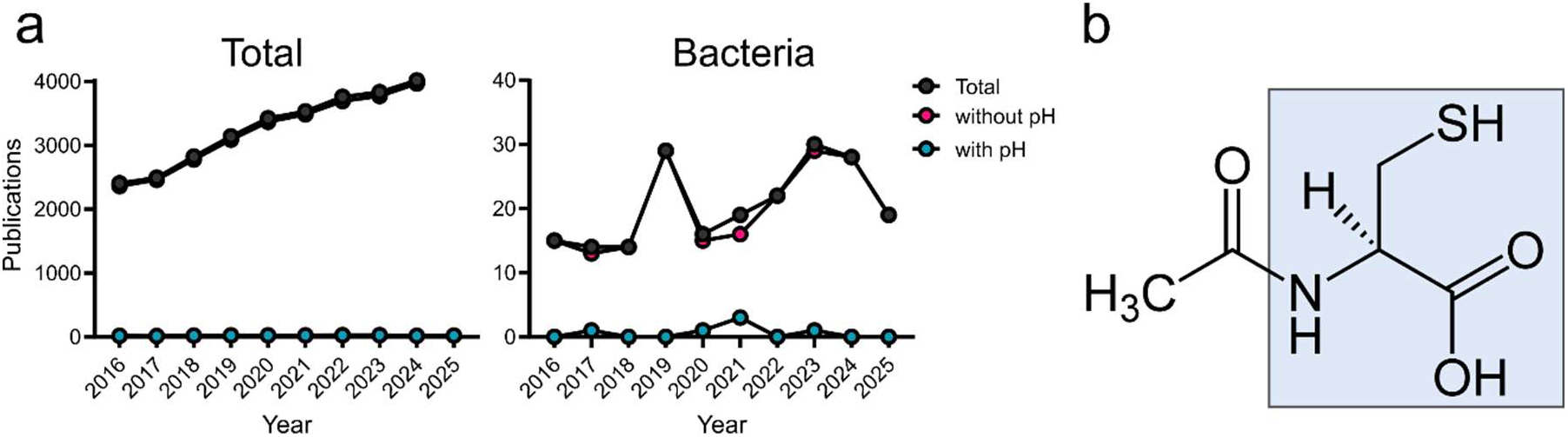
(a) Metrics of yearly publications from 2016-2025 mentioning NAC in total or with bacteria, divided into total number of publications, pH not mentioned, or mentioned in title or abstract. Raw data obtained from dimensions.ai using the search terms NAC / N-acetylcysteine / acetylcysteine AND bacteria* and NOT pH or AND pH for differentiation. (b) Chemical formular of N-acetylcysteine with cysteine highlighted in blue.

NAC is composed of the sulfur-containing amino acid cysteine with an acetyl group attached to its nitrogen atom (Fig. 1b). Compared with cysteine, NAC has higher water solubility in its monomeric (100 mg/mL) and oxidized (1000 mg/mL) forms (15, 16), and a higher protonation constant (pKa) of its thiol group (9.51 versus 8.18), reducing its reactivity at physiological pH which contributes to higher stability in solution (16). Despite this increased stability, NAC is still prone to oxidation, forming dimers (17, 18), which is reported to hinder its anti-oxidative ability (19) and conversion to cysteine (18).

The carboxylic acid (-COOH) and sulfur (-SH) groups give NAC two pKa values, which specifies the dissociation of hydrogen from the functional groups: pKa of 9.51 for -SH and pKa of 3.24 for -COOH (20). According to the Henderson-Hasselbach equation, at pH=pKa, 50% of the weak acid is ionized, and ionization increases logarithmically with further increasing pH, making most molecules deprotonated and giving NAC a negative charge at neutral pH – in contrast to cysteine. Since only uncharged molecules readily diffuse across biological membranes (21), NAC ionization state is expected to directly influence antibacterial activity.

The antibacterial mechanism of NAC related to its weak acid properties has been systematically investigated primarily in the gram-negative bacterium *P. aeruginosa*. Kundakad *et al.* compared the actions of NAC to acetic acid, another weak organic acid, and demonstrated that both compounds required a pH below their pKa values to exert antibacterial and antibiofilm activity. They proposed that non-ionized NAC diffuses into bacterial cells, where intracellular deprotonation leads to s acidification and bacterial killing. In line with this, NAC at pH 3.3 killed *P. aeruginosa* at 10 mg/mL (approximately 62 mM), whereas 60 mg/mL at neutral pH had no antibacterial effect. (22). This pH-dependent mechanism was later confirmed in wound-derived *P. aeruginosa* isolates (23). However, to our knowledge, no systematic mechanistic study has examined whether similar pH-dependent antibacterial effects occur in the gram-positive pathogen *S. aureus*.

The few studies assessing pH-dependent effects on *S. aureus* use slightly different setups and report different outcomes. Supporting the hypothesis that a pH<pKa is necessary, Kunkukad *et al.* varied pH by using LB or PBS as solvent and found NAC to be bactericidal at 2 mg/mL at pH 3, but inactive at pH 3.8 (24). In contrast, several studies reported antibacterial effects of pH-neutral NAC against *S. aureus,* including MICs ranging from 24 mg/mL (25) to 128 mg/mL (9), bactericidal activity at 20-200 mM (26) and bacteriostatic effect with enhanced regrowth at 30 mM (12).

The established pH dependence of NAC antibacterial activity in *P. aeruginosa*, together with the conflicting findings reported for *S. aureus*, prompted us to systematically investigate the pH dependence of NAC activity against *S. aureus*. In addition, because sublethal concentrations of antimicrobial compounds can alter bacterial physiology in ways relevant to virulence and antibiotic tolerance, we further investigated whether pH-neutral NAC perturbs bacterial metabolism under both planktonic and biofilm growth conditions.

## Results

### Antibacterial activity of NAC is pH-dependent

Planktonic growth of the clinical *S. aureus* strain Mu12 was assessed in TSB supplemented with 0–200 mM NAC, either adjusted to pH 7 or without pH-adjustment. At neutral pH, NAC did not affect bacterial growth across the tested concentration range (Fig. 2a). This lack of growth inhibition persisted even at concentrations approaching the NAC solubility limit of 100 mg/mL (80 mg/mL tested, Fig. S1). In contrast, non-adjusted NAC reduced growth by approximately 22% at 20 mM (pH 6.3) and completely inhibited detectable growth at 200 mM (pH 2.8) (Fig. 2b). To distinguish bacteriostatic from bactericidal effects, bacteria exposed to NAC for 24 h were transferred to fresh TSB and assessed for regrowth. CFU enumeration showed no differences between conditions with detectable growth after NAC exposure, whereas no viable bacteria were recovered following treatment with 200 mM non-adjusted NAC, confirming bactericidal activity under acidic conditions (Fig. 2c). Similar pH-dependent effects were observed in the *S. aureus* USA300 strain (Fig. S2), indicating that the observed effects were not strain-specific.

**Fig. 2.**
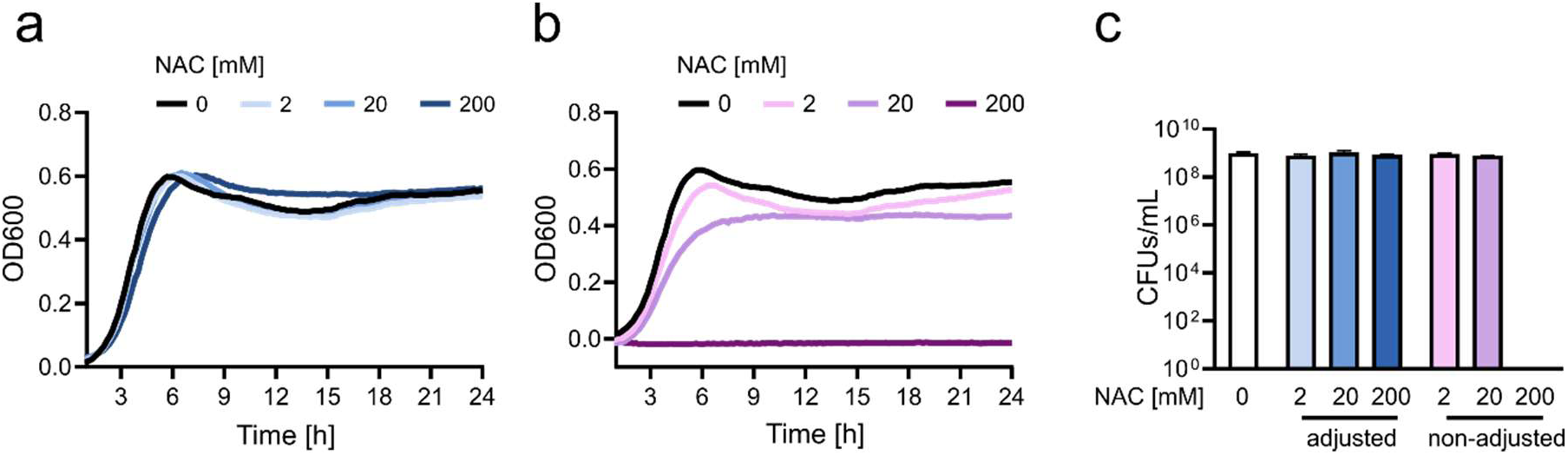
Antibacterial activity of NAC against *S. aureus* Mu12 is pH-dependent. Growth curves obtained by OD_600_ measurements over 24 hours in TSB supplemented with 0, 2, 20, or 200 mM of NAC (a) with prior adjustment to pH 7 or (b) without pH adjustment. Curves represent the mean of three independent experiments. (c) Bacterial re-growth assessed by over-night CFU count of bacteria following transfer to fresh TSB after 24-hour exposure to 0, 2, 20, or 200 mM NAC with or without pH-adjustment to 7. Data represent mean ± standard deviation (SD) of three technical replicates.

We next investigated whether pH adjustment with NaOH altered NAC stability and thereby contributed to the observed loss of antibacterial activity at neutral pH. Quantification of NAC concentrations showed no differences over time between pH-adjusted and non-adjusted 200 or 2 mM NAC solutions (Fig. 3a). Since NAC concentrations gradually decreased in both 2 mM solutions, potential degradation products including cysteine (deacetylation), N-acetylalanine (desulfurization), and alanine (deacetylation and desulfurization) were additionally quantified. None differed across conditions (Fig. S3a), indicating that no measurable NAC degradation occurred under the tested conditions. In contrast, NAC dimerization increased over time in all groups (Fig. S3b), leading to increasing diNAC/NAC ratios (Fig. 3b). Dimerization ratios were significantly higher in pH-adjusted 200 mM solutions, where the pH difference between conditions was the highest, whereas no differences were observed at 2 mM (Fig. S3c). These findings are consistent with reduced oxidative dimerization of the NAC thiol group under acidic conditions.

**Fig. 3.**
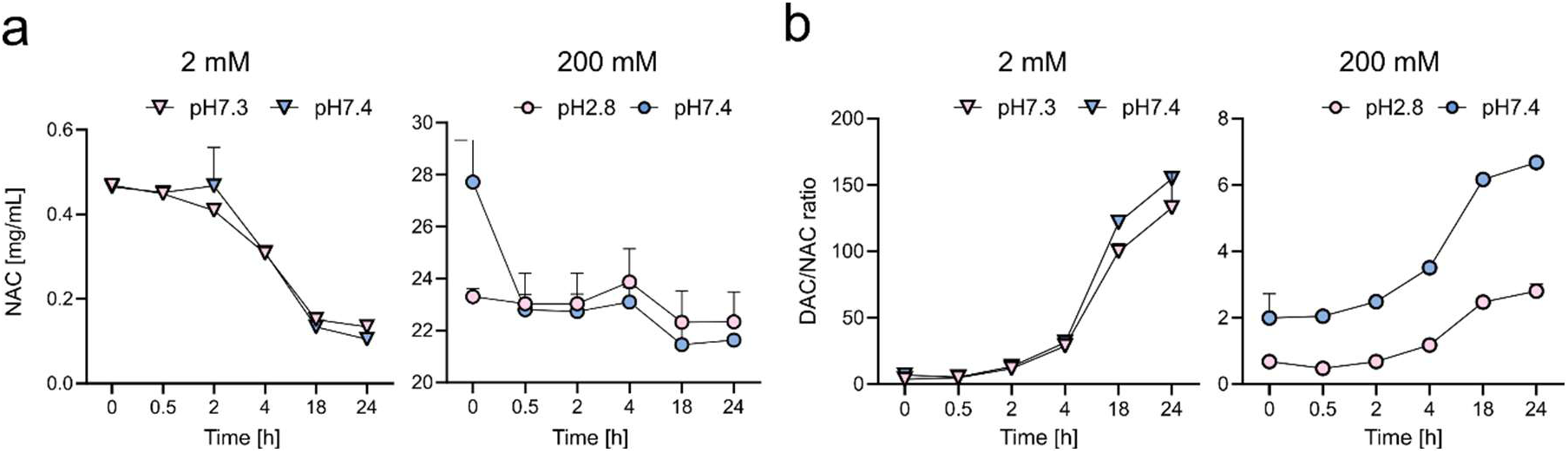
Neutralization of pH increases NAC dimerization ratio. Sterile 2 and 200 mM NAC solutions with and without pH-adjustment were incubated in TSB at 37°C for up to 24 hours to quantify NAC and its degradation products. (a) Calculated NAC content (mg/mL), and (b) dimer to monomer (diNAC/NAC). Data represent mean ± SD of three replicates.

### pH-neutral NAC impacts planktonic *S. aureus* metabolism in a concentration-dependent manner

Because pH-neutral NAC (pH 7.4) did not inhibit growth of *S. aureus*, we next investigated whether it impacts bacterial metabolism, as NAC serves as a direct precursor of the sulfur containing amino acid cysteine. Intra- and extracellular metabolites were quantified in late stationary-phase planktonic Mu12 bacterial cultures. Principal component analysis (PCA) showed a clear separation of intracellular metabolite profiles along PC1 (Fig. 4a), indicating a global metabolic response to NAC. In line with this, numerous intracellular metabolites correlated with NAC concentration, a complete list is provided in Table S.1.

**Fig. 4.**
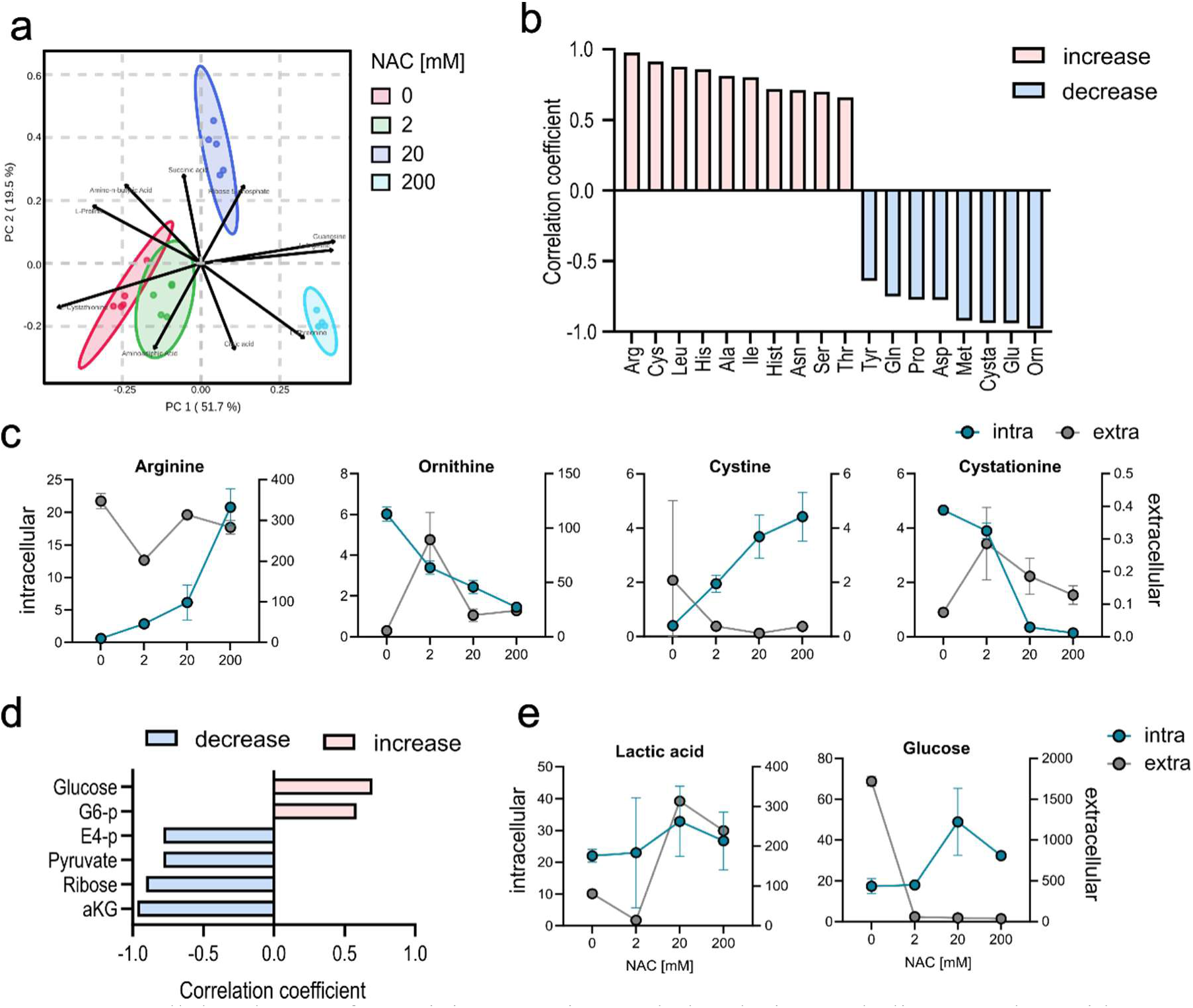
Intracellular changes for arginine, cysteine, and glycolysis metabolites correlate with NAC supplementation in planktonic stationary-phase Mu12. (a) PCA plot of intracellular metabolite abundances. (b) Significant correlations between intracellular amino acid abundances and supplemented NAC concentrations. (c) Quantified concentrations of selected arginine and cystine metabolites expressed in μM. Intracellular metabolites plotted on left and extracellular metabolites on the right y-axis. (d) Significant correlations between intracellular glucose and TCA metabolites and supplemented NAC concentrations. (e) Quantified concentrations of lactic acid and glucose expressed in μM, Intracellular metabolites plotted on left and extracellular metabolites on the right y-axis. Correlation was determined using Pearson correlation with a threshold of FDR <0.05. Five replicates per group were analyzed.

The strongest intracellular correlations with NAC concentration were observed in amino acid metabolism (Fig. 4b), namely in arginine and cysteine metabolism. Arginine itself showed the strongest positive correlation, while its precursors/degradation products ornithine and glutamate showed the strongest negative correlation. These changes in arginine and its precursors were not mirrored extracellularly (Fig. 4c), suggesting intracellular metabolic reprogramming rather than altered uptake. In cysteine metabolism, the NAC derived cysteine dimer cystine increased with NAC concentrations, whereas its connected metabolites cystathionine and methionine decreased. As observed for arginine metabolism, these trends were restricted to intracellular metabolites.

Metabolites associated with glycolysis and TCA cycle were also affected by NAC supplementation. The TCA cycle intermediate α-ketoglutarate decreased with increasing NAC concentration, whereas glucose and its downstream metabolite glucose 6-phosphate increased, and the glycolysis end product pyruvate decreased (Fig. 4d). These intracellular changes were accompanied by reduced extracellular glucose and increased lactic acid levels (Fig. 4e), indicating enhanced glycolytic flux with NAC addition.

### pH-neutral NAC induces similar concentration-dependent changes in biofilm *S. aureus* metabolite abundances

We next examined whether the metabolic changes observed in planktonic bacteria were conserved under biofilm-growth conditions, given that NAC is often used as an anti-biofilm agent and that bacteria within biofilms exhibit a distinct metabolic state. As observed for planktonic cultures, neither CFU counts nor metabolite medians differed significantly across conditions (Fig. S4). Although NAC-induced intracellular correlations in biofilms were fewer in number and weaker in correlation strength, 9 out of 15 positive and 6 out of 12 negative correlations overlap with planktonic changes (Fig. 5a). The affected pathways again included amino acid metabolism and central carbon metabolism (Table S.2).

**Fig. 5.**
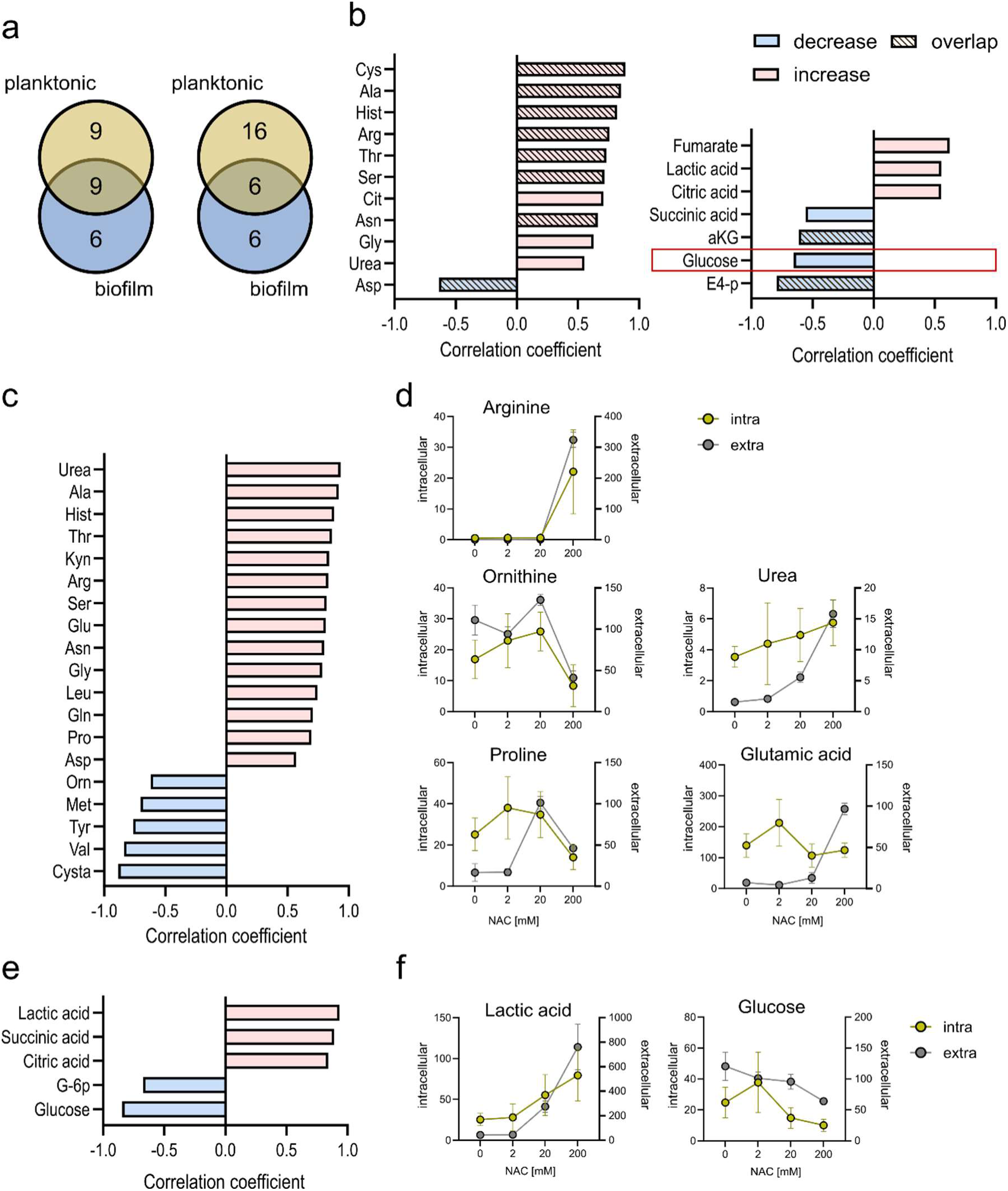
Metabolite abundances for energy metabolism, and cystine and arginine metabolism correlate with NAC concentration in 3-day old biofilm Mu12. (a) Overlaps of metabolites correlating with supplemented NAC concentration between growth modes, separated in positive (left) and negative (right) correlations. (b) Significant correlations between intracellular amino acid (left) and energy metabolites (right) with supplemented NAC concentration. Overlaps with planktonic correlations as shaded bars, reversed correlation indicated with red outline. (c) Significant correlations between extracellular amino acids and NAC concentrations. (d) Quantified intra- and extracellular concentrations of metabolites connected to arginine metabolism expressed in μM. Intra- and extracellular concentrations plotted on left and right y-axis respectively. (e) Significant correlations of extracellular glycolysis and TCA cycle metabolites with NAC molarity. (f) Quantified intra- and extracellular concentrations of glycolysis metabolites expressed in μM. Intra- and extracellular concentrations plotted on left and right y-axis respectively. Significance threshold for correlation at an FDR<0.05. Five replicates per group were analyzed.

Key overlapping intracellular correlations between growth conditions included increased cystine and arginine levels. Biofilm-specific changes included the arginine precursor citrulline and the degradation product urea, both of which increased with NAC concentration (Fig. 5b). Among metabolites associated with glycolysis and TCA cycle, only decreasing α-ketoglutarate was shared with planktonic cultures, whereas increased lactic acid and decreased glucose were biofilm specific. Notably, glucose was the only metabolite exhibiting opposite correlation patterns between growth conditions, showing positive correlations with NAC concentration in biofilm bacteria and negative correlations in planktonic cultures.

In biofilm bacteria, intracellular correlations were accompanied by numerous extracellular correlations with NAC concentrations (Fig. 5c). As observed for intracellular metabolites, NAC had the highest effect on amino acid concentrations. Strong increases were found for arginine itself, its precursors glutamic acid and proline, and degradation product urea, whereas ornithine decreased. These changes occurred simultaneously intra- and extracellularly (Fig. 5d), suggesting altered arginine metabolism rather than simply increased import.

In contrast to planktonic cultures, intracellular cystathionine and the downstream metabolite methionine did not decrease in biofilm bacteria. However, both metabolites decreased in biofilm supernatants, pointing towards altered transport or consumption dynamics in 3-day-old biofilms. For central carbon metabolism, decreasing glucose and glucose 6-phosphate paired with marked lactic acid increase (Fig. 5e), which again occurred both intra- and extracellularly (Fig. 5f). The simultaneous decreases of glucose intra- and extracellularly and increases of lactic acid suggest an increase of glycolytic flux accompanied by enhanced glucose deprivation.

### Growth-mode specific metabolism is preserved despite NAC-induced metabolic changes

The overlaps between NAC-responsive pathways in planktonic and biofilm bacteria, prompted to compare the relative metabolite abundances across both growth conditions. To establish a baseline for this comparison, planktonic and biofilm bacteria without NAC supplementation were first compared. Biofilm growth was associated with substantial alterations in arginine, cysteine, and TCA and glycolysis metabolism (Table 1), mirroring the pathways affected by NAC supplementation.

**Table 1.**
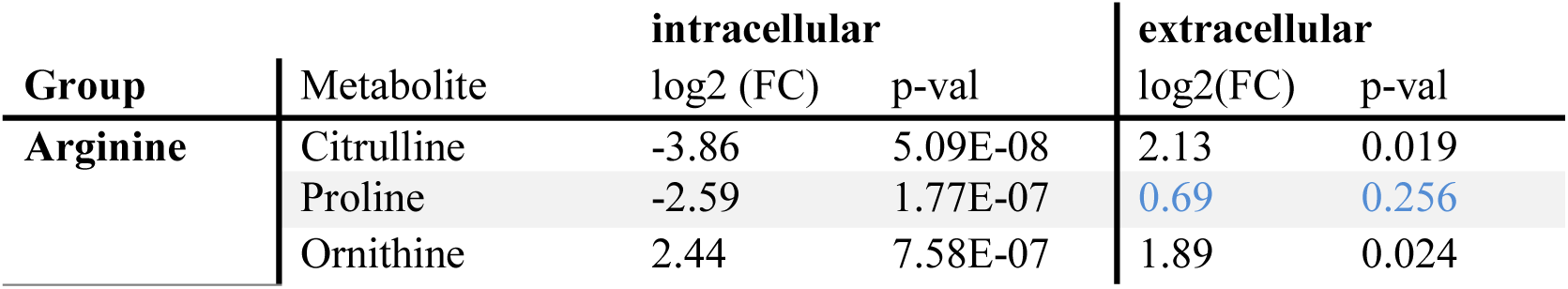
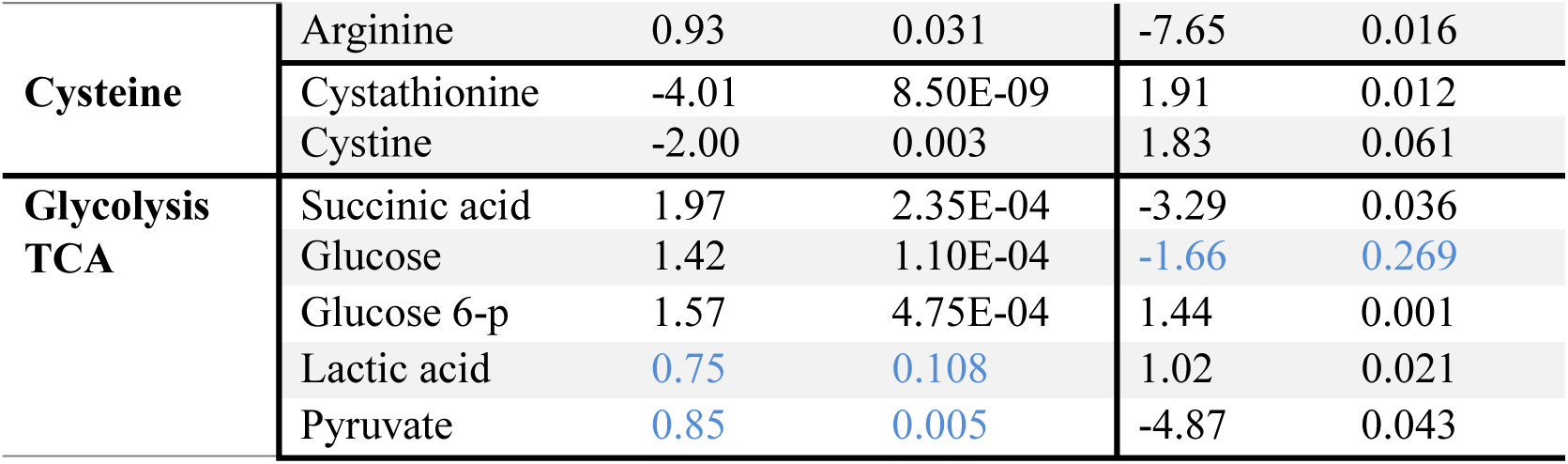
Significantly changed metabolites in biofilm bacteria belonging to arginine, cysteine and glycolysis, and TCA cycle. Significant values in black, non-significant values in light blue for completeness. Exploratory significance threshold of FC>2 and p<0.1 used.

Compared with planktonic cultures, bacteria within the biofilm exhibited strong alteration of arginine metabolites, with decreased intracellular citrulline and proline, increased ornithine, and strong extracellular arginine depletion, indicating increased arginine consumption and catabolism. Cysteine metabolism was similarly altered, with decreased intracellular cystine and cystathionine. In central carbon metabolism, intracellular glucose and succinic acid increased, while extracellular lactate accumulated. Changes in energy metabolites suggests increased glycolysis and lactic acid fermentation and decreased TCA cycle activity in biofilm bacteria (Table 1).

Although the metabolic pathways altered between growth conditions overlapped with NAC-induced changes, the distinct overall metabolic profiles of planktonic and biofilm bacteria were not disturbed following NAC supplementation (Fig. 6a). PCA showed clear separation of planktonic and biofilm samples along PC1 (45.5% explained variability), whereas sample distribution along PC2 (24% explained variance) corresponded to NAC concentrations, underlining the previous conclusion that NAC induced metabolic changes are mirrored in both growth condition.

**Fig. 6.**
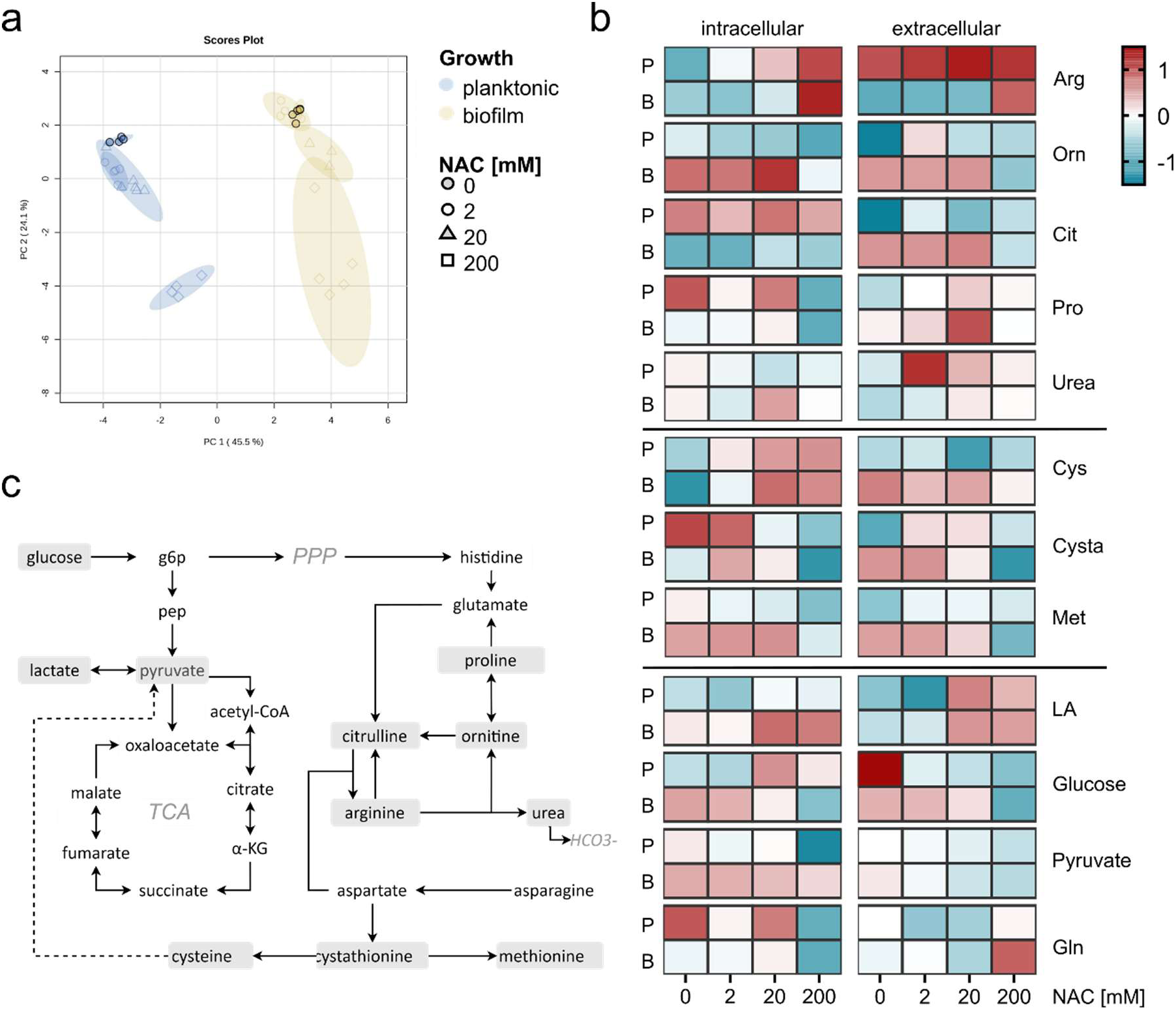
NAC induces overlapping metabolic shifts in planktonic and biofilm Mu12 without overriding growth condition-specific metabolic profiles (a) PCA plot of relative metabolite abundances in planktonic and biofilm bacteria with and without NAC supplementation. (b) Heatmap visualization of metabolites connected to arginine (top), cysteine (middle), and glycolysis and TCA cycle metabolites (bottom) in planktonic and biofilm samples. Planktonic (P) and biofilm (B) conditions are shown for each metabolite, left portion displays intracellular and right portion extracellular metabolite abundances. (c) Schematic overview of measured metabolites and pathway connections. Grey highlights denote metabolites included in the heatmap visualization. Pathways are based on KEGG annotations for USA300, with additional connections indicated by dashed lines.

Since arginine, cystine, and TCA and glycolysis metabolites were impacted by both NAC supplementation and bacterial growth condition, these pathways were further examined on the intra- and extracellular changes (Fig. 6b) and their connections (Fig. 6c). Differences in arginine-associated metabolites between planktonic and biofilm bacteria became less pronounced with increasing NAC concentration, accompanied by increased arginine abundance under both growth conditions. This data suggests that biofilms consumed more arginine at baseline, whereas NAC addition induced arginine accumulation.

The cysteine precursor NAC reversed intracellular cystine depletion in biofilm bacteria, leading to intracellular cystine accumulation under both growth conditions. This was accompanied by decreasing cystathionine levels both intra- and extracellularly under both growth conditions, and a similar decrease in methionine levels, although the magnitude of these changes differed between biofilm and planktonic conditions. (Fig. 6b)

Growth attributed differences in glycolysis did not become less pronounce with NAC addition. Lactic acid accumulation both intra- and extracellular characterized biofilm bacteria and continued increasing for both growth forms with increasing NAC addition. Glucose, which was initially depleted extracellularly for biofilm bacteria, reduces for both growth conditions with NAC addition. The same dynamic is mirrored by pyruvate (Fig. 6b). Taken together, this suggests that glycolysis and lactic acid fermentation increased under biofilm conditions, and NAC addition further increases glucose uptake as well as lactic acid fermentation in both growth conditions.

## Discussion

This study demonstrates that the antibacterial activity of NAC against *S. aureus* is strongly pH-dependent and is abolished following pH neutralization. In contrast, pH-neutral NAC did not inhibit bacterial growth but induced substantial metabolic alterations under both planktonic and biofilm growth conditions. These findings help reconcile highly inconsistent reports of NAC antibacterial activity in the literature and indicate that NAC can alter bacterial physiology independently of direct growth inhibition.

In this study we report that NAC is bactericidal against *S. aureus* at a pH<pKa, an activity that ceases completely when pH was neutralized. Both tested strains had reduced growth at 20 mM (3.26 mg/mL) and were killed by 200 mM (32.6 mg/mL) NAC without pH-adjustment (pH 2.8). This concentration is within the upper bounds of previous reports for clinical *S. aureus* isolates, which ranges from 2.5 mg/mL (8) up to 32 mg/mL (26). The pH-dependent antibacterial effects found by Kundukad *et al.* (24) were in accordance with our findings, supporting that a pH below pKa is likely the necessary condition for NAC’s antibacterial activity.

In addition, we found that pH-adjustment increased dimerization but did not cause measurable degradation at 37°C over a 24-hour period. NAC dimerization was significantly lower at low pH. In general, NAC degradation is mostly referenced as decreasing NAC content (27, 28), or as oxidation to its dimeric form diNAC (18, 19). The dimerization at 37°C is supported by previous reports (17). The lack of significant NAC decrease at high concentrations supports that NAC’s pH-dependent effects were not due to degradation but most likely due to protonation.

Although NAC at neutral pH had no effect on growth, we quantified metabolic changes because NAC is being investigated as an antibiotic adjuvant (12–14) and metabolism appears to be linked to antibiotic resistance and virulence – as discussed in the following sections. We found systematic and concentration-dependent shifts in metabolite abundances and highlighted the affected pathways of arginine and cysteine metabolism, glycolysis and TCA cycle as their changes partially overlap between the growth modes of the clinical isolate Mu12.

As NAC is a direct precursor of cysteine (29), the observed concentration-depended accumulation of intracellular cystine is consistent with cellular uptake and conversion of NAC. Whether the anionic form of NAC itself is transported into *S. aureus* or first converted to cysteine prior to uptake cannot be conclusively stated from our results. That said, although the known sulfur acquisition transporters TcyABC and TcP prefer cystine for transportation, they are able to use both cystine and NAC as substrates (30, 31), suggesting that the metabolic perturbations observed following NAC addition may involve bacterial uptake of NAC.

Regardless of the import route, cysteine accumulation itself has downstream consequences. It represses the cysteine master regulator CymR, which in turn represses toxin production and promotes methionine utilization (30, 32). The observed negative correlation of cystathionine and methionine with NAC concentration under both conditions is consistent with CymR-mediated suppression of methionine utilization pathways. Interestingly, intracellular accumulation of cysteine has been linked with medium acidification (32), likely through its conversion to pyruvate (32, 33) – a connection that may link the cysteine finding to the glycolysis as well as arginine changes discussed below.

NAC supplementation was associated with increasing glucose uptake and lactic acid accumulation under both planktonic and biofilm conditions, suggesting an increasing disconnection between glycolysis and TCA cycle activity. Generally, *S. aureus* TCA cycle activity is repressed under nutrient-rich conditions, aiding rapid cellular division and diversion of carbon to fermentation and the pentose phosphate pathway (34). TCA cycle activity is resumed during the post-exponential phase due to the limited availability of preferred carbon sources (35) – a condition present in both growth conditions we analyzed. When TCA cycle flux is reduced, pyruvate is redirected toward fermentation, producing acetate and lactic acid rather than entering the TCA cycle (34, 36). Our findings are consistent with this model: increasing NAC concentrations appear to deepen this bottleneck. However, we cannot conclusively distinguish increased fermentation from impaired lactic acid recycling to pyruvate via lactate dehydrogenase (37), as conversion of cysteine to pyruvate may impair this mechanism.

Our observation of elevated lactic acid fermentation under biofilm conditions is well documented, possibly driven by reduced oxygen tension within the biofilm matrix (37, 38). The observation that NAC further increases lactic acid production under biofilm conditions suggests it amplifies an existing metabolic shift rather than introducing a new one. This is consistent with the partial but not complete overlap of metabolite correlations between the two growth modes: NAC pushes both toward a shared metabolic state, but does not completely override the fundamental differences imposed by biofilm physiology. The connection between energy metabolism and biofilm formation is complex – although lactic acid production is increased, TCA components are also enriched in biofilms at the transcription and protein level analysis (39–41), yet forced TCA cycle induction reduces biofilm formation (42), and glycolysis-TCA cycle coupling was found to be disrupted in biofilm bacteria (43). Our data do not suggest that NAC addition disrupts biofilm energy metabolism.

The most striking metabolic response to NAC addition was the strong accumulation of arginine both intra- and extracellularly, accompanied by changes in its precursors and degradation products that together suggest increased arginine biosynthesis. Despite encoding two complete arginine synthesis pathways – a canonical glutamate-to-arginine route and an *S. aureus*-specific proline-to-arginine route (33, 44) – *S. aureus* is classically considered arginine auxotrophic (44–46), because of synthesis repression (47). This assumption, however, is based on laboratory strains. During infection, arginine prototrophs were found to emerge as the dominant population, and 50% of clinical isolates seem to be able to satisfy arginine requirements through proline. (46, 48) The clinical isolate Mu12 used in this study may therefore be capable of de novo arginine synthesis, and NAC addition appears to trigger or enhance this capacity.

The biological significance of the observed arginine accumulation is underscored by arginine’s established roles in *S. aureus*. Increased arginine metabolism and urea cycle upregulation have been documented in biofilm bacteria and interpreted as a response to fermentation-driven acidification (39, 41, 49). At baseline in our data, arginine was increasingly consumed under biofilm conditions – consistent with the reports in the literature. This consumption was reversed by NAC addition, suggesting that NAC-driven arginine synthesis may contribute to pH homeostasis. Consistent with this model, we found coinciding increases of lactic acid and arginine in intra- and extracellular compartments. In the context of infection, arginine metabolism has been linked to antibiotic tolerance, where both restriction, and increased degradation paired with decreased synthesis promoted tolerance *in vitro* (50), while arginine supplementation *in vivo* attenuated infection severity (51). Whether NAC-induced arginine accumulation influences antibiotic susceptibility in Mu12 is an open question warranting further investigation.

In conclusion, we demonstrated that NAC’s antibacterial properties against *S. aureus* are pH dependent. Despite a lack of effect on bacterial growth, our findings suggest that NAC is imported into the bacteria, leading to cysteine accumulation, combined with increased glycolysis, lactic acid accumulated both intracellularly and extracellularly. These changes triggered increased arginine synthesis, most likely to counteract extracellular acidification caused by NAC addition. Changes in energy metabolism did not suggest that NAC hinders biofilm metabolism or aids its destruction. However, the strong induction of arginine synthesis caused by NAC warrants further investigation into how NAC and the metabolic disturbances caused can aid antibiotic properties and how the triggered metabolic disturbances impact bacterial virulence.

### Conclusions

Our results demonstrate that the antibacterial activity of NAC against *S. aureus* is primarily driven by pH rather than the compound itself. Although pH-neutral NAC does not impair growth, it induces concentration-dependent metabolic changes in both planktonic and biofilm bacteria. A consistent increase in intracellular arginine and cystine, alongside enhanced glycolytic flux and lactate production, suggests a shift toward altered redox balance and energy metabolism in response to NAC. These findings indicate that NAC modulates bacterial physiology without directly inhibiting growth. Understanding these metabolic adaptations provides a basis for exploring NAC as a modulator of bacterial behavior rather than solely as a classical antimicrobial agent.

## Materials and Methods

### Materials and reagents

Reagents for bacterial culture were obtained from Thermo Fisher scientific: Tryptic soy broth (TSB, CM029B); phosphate buffered saline (PBS, P4417) tablets, N-acetylcysteine (NAC, A7259-25G), sodium hydroxide (NaOH, 71690) for pH-adjustment; and Liofilchem: tryptic soy agar plates (TSA, 10037). Reagents for LC-MS measurements were obtained from Thermo Fisher scientific: ammonium bicarbonate (A6141); HPLC grade methanol (34885-M).

### Bacterial strains

Two *S. aureus* strains were used: USA300 obtained from ATCC, and the clinical isolate Mu12 obtained from an FRI case the Unfallklinik Murnau, Bavaria, Germany. Identification and antibiogram results can be found in Table S.3). All bacteria were stored cryopreserved in TSB with 80% glycerol at −20°C.

### Preparation of reagents

TSB for inoculum preparation was autoclaved at 121°C for 20 minutes; TSB for experiments was prepared as 1x and 2x solution and filter sterilized with a 0.22 μm filter (TRP, 99722) to unify conditions across dilutions.

NAC working solutions for growth experiments and LC-MS measurement were prepared with and without pH-adjustment as follows: Non-adjusted stocks were prepared as 100 mg/mL NAC in deionized water. Adjusted stocks were prepared as 100 mg/mL NAC in 0.609 M NaOH solution in deionized water. All stocks were incubated in a 37°C water bath for 10 minutes to aid dissolution. Before dilution of NAC stocks to 400 mM with deionized water, pH was measured (Metrohm, 827 pH lab) and pH-corrected stocks were adjusted to pH 7 (±0.05) by dropwise addition of 1 M NaOH, to increase precision across experiments with varying total volumes.

These 2x stock solutions were filter sterilized as described above for TSB preparation and combined 1:1 (v/v) with 2x TSB stocks to obtain final working solutions with 200 mM NAC and further diluted with filter sterilized 1x TSB to obtain 20 and 2 mM working solutions. The pH was recorded for all solutions, summarized in Table 2. Only 200 mM NAC in TSB reached a pH < NAC’s pKa of 3.24.

**Table 2.**
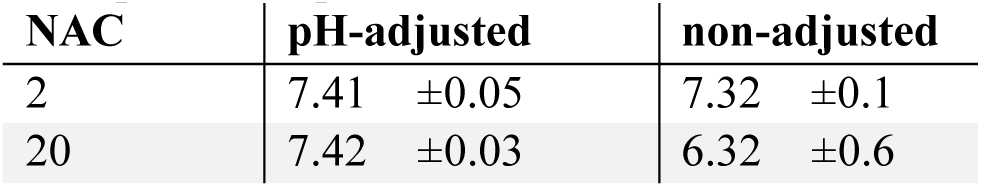
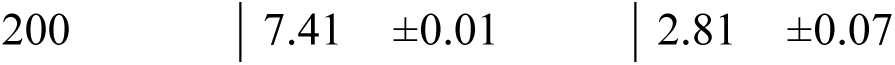
pH measurement of NAC containing medium expressed as median ± standard deviation, 3 independent experiments used.

| NAC | pH-adjusted | non-adjusted |
| --- | --- | --- |
| 2 | 7.41 $\pm 0.05$ | 7.32 $\pm 0.1$ |
| 20 | 7.42 $\pm 0.03$ | 6.32 $\pm 0.6$ |

### Inoculum preparation

All bacteria tested were inoculated in TSB and grown over night at 37°C under constant agitation (180 rpm). Before seeding, over-night cultures were diluted 1:20 in fresh TSB and cultured for 2 hours to regain log growth and washed twice with PBS by pelleting at 1320 rcf for 5 minutes and resuspending in PBS. After washing, OD_600_ was measured (MultiskanGO, Thermo Scientific) and adjusted with PBS to 0.6 for planktonic experiments and to 1 for biofilm experiments.

### Determining the effects of NAC on planktonic growth

Growth curves were recorded in 96 well plates. For this, 200 μL of adjusted or non-adjusted NAC solutions were added in 6 replicates and washed bacterial suspensions were added to a final OD_600_ of 0.01. Not inoculated medium was used as negative controls in triplicates. Bacteria were grown at 37°C under constant agitation (180 rpm) for 24 hours, OD_600_ was measured every 10 minutes using a microplate reader (Labrador, Cerillo).

Re-growth of bacteria after 24-hour incubation with NAC was tested to differentiate bacteriostatic and bactericidal effects. Bacteria were grown for 24 hours in 0-200 mM NAC with or without pH-adjustment as described for growth curves. After incubation, 20 μL of bacterial suspension was transferred to a 96 well plate containing 180 μL of fresh TSB and bacteria were incubated overnight at 37°C under constant agitation (180 rpm). Bacterial suspensions were serial diluted in sterile PBS, plated on TSA plates, and incubated overnight at 37°C to count colony forming units (CFUs) (Interscience Scan 1200).

To test if growth inhibition occurs at higher concentrations of pH-neutral NAC, highly concentrated working solutions were obtained by using pH-adjusted NAC stock to solve TSB powder directly. This way we obtained NAC solutions of 80 mg/mL (490 mM) - close to NAC’s solubility maximum of 100 mg/mL. Medium was filter sterilized as described above. 400 μL were added to 96 well plates and 200 μL were used for 1:1 (v/v) serial dilution with filter sterilized TSB. Serial diluted plates were inoculated with bacterial suspension to a final OD_600_ of 0.01 and growth curves determined with OD_600_ measurement as described above. All growth experiments were conducted with both Mu12 and USA300 strains.

### Investigating the effects of NAC on planktonic metabolism

Growth media was prepared from 200 mM pH-adjusted NAC solutions as described in the growth section, and filter sterilized TSB used as 0 mM control. For planktonic cultures, 20 mL of medium supplemented 0, 2, 20, or 200 mM NAC were added to Erlenmeyer flasks and bacterial inoculum was added to a final OD_600_ of 0.01. Bacteria were grownat 37°C under constant agitation (150 rpm) for 16 hours.

### Investigating the effects of NAC on biofilm bacterial growth and metabolism

For biofilm cultures, bacterial cultures and pH-adjusted NAC containing media were prepared as described above, and inoculum was added to 6 mL prepared medium at a final OD_600_ of 0.1. Steam sterilized titanium disks (inhouse manufactured) were placed in 24 well plates, and 1 mL inoculated medium was added in 5 replicates per group. Non-inoculated medium with 0 and 200 mM were used as negative controls. Biofilms were cultured at 37°C under constant agitation (20 rpm) for a total of three days. Medium was changed every 24 hours by replacing the entire volume with freshly prepared medium.

Biofilm bacterial CFU count was determined to make sure that growth differences do not affect metabolomic comparison. For this, titanium disks were transferred to a new plate and washed twice with 1 mL PBS. After covering with 1 mL PBS, disks were sonicated for 3 minutes to disrupt the biofilm and 150 μL suspension were serial diluted in sterile PBS and plated on TSA plates. Plates were incubated overnight at 37°C and colonies counted (Interscience Scan 1200).

### Planktonic and biofilm culture harvest for metabolomics measurement

Planktonic cultures were harvested for extra- and intracellular metabolites. 50 μL of suspension was collected in triplicates from the growth flask, centrifuged at 21,130 rcf, 4°C for 3 minutes. Supernatants were collected and frozen at −80°C for analysis of extracellular metabolites. To obtain bacteria for intracellular harvesting, OD_600_ of bacterial suspension was measured and adjusted volumes were collected to reach a final bacterial count of approximately 5 x 10^8^ cells per tube. After harvest, bacteria were pelleted at 10,000 rcf, 4°C for 3 minutes, resuspended in 1 mL ice-cold 10 mM ammonium bicarbonate (Sigma Aldrich, A6141) and pelleted again at 10,000 rcf, 4°C for 3 minutes. Tubes were immediately put on an ice block, bacteria resuspended in 1.8 mL cold 100% methanol (Sigma Aldrich, 34885-M), and snap frozen in liquid nitrogen. Samples were stored at −80°C until further processing.

Biofilm cultures were harvested for extra- and intracellular metabolites. 50 μL of suspension was collected from each well, centrifuged at 21,130 rcf, 4°C for 3 minutes. Supernatants were collected and frozen at −80°C for analysis of extracellular metabolites. Biofilms were washed once with 1 mL ice cold 1 mM ammonium bicarbonate, and titanium disks transferred to a fresh plate on ice containing 1 mL cold 100% methanol. Bacteria were harvested by scraping. After transferring the liquid, disks were washed once with 1 mL ice cold methanol, pooled samples were snap frozen in liquid nitrogen, and samples were stored at −80°C until further processing.

Intracellular metabolites for both harvests were extracted using freeze-thaw method based on (52). In brief, bacteria were freeze-thawed three times by thawing at room temperature for 7 minutes and freezing in liquid nitrogen for 3 minutes. After vortexing for 5 seconds, lysates were centrifuged at 21,130 rcf at 4°C for 4 minutes. The obtained supernatants were collected and stored at −80°C until final preparation.

### Targeted LC-MS measurement of metabolites

We quantified intra- and extracellular metabolites using LC-MS based targeted metabolomics approach. The measured metabolites included central carbon metabolism (glycolysis, TCA cycle, pentose phosphate pathway), amino acids and derivatives, (poly)amine metabolism, and fatty acid oxidation.

For bacteria culture supernatants, 10 μL of supernatant was mixed with 80 μL methanol and 10 μL isotopically labelled internal standard. Samples were vortexed briefly, centrifuged at 17,850 rcf for 5 minutes, and 90 μL of supernatant was transferred to HPLC vials for measurement. Intracellular extracts were centrifuged at 17,850 rcf for 5 minutes and supernatants dried completely in a vacuum centrifuge (Concentrator plus, Eppendorf) on V-AL for approximately 3 hours. Dried extracts were reconstituted in 90 μL methanol and 10 μL isotopically labeled internal standard, vortexed briefly, centrifuged at 17,850 rcf for 4 minutes, and supernatants transferred to HPLC vials for measurement.

Targeted quantitative metabolite analysis was conducted using HILIC-based liquid chromatography and mass spectrometric detection. Metabolites were separated on ACQUITY Premiere BEH Z-HILIC analytical columns (1.7 μm 2.1 x 100 mm, Waters) using a gradient elution method with mobile phase A (0.15% formic acid, 10 mM ammonium formate in water) and mobile phase B (0.15% formic acid, 10 mM ammonium formate in 85% acetonitrile), with a total analysis time of 18 minutes. The mobile phase flow rate was set to 0.4 mL/minute, injection volume to 2 μL, and column temperature to 40 °C.

Metabolites were detected on an Orbitrap Exploris 120 (Thermo Fisher Scientific) using ESI positive and negative modes in full scan detection. Scan range was set to 100 - 500 m/z, and mass resolution to 30000. ESI spray voltage was set to 3.5 kV (positive mode) and 2.5 kV (negative mode), gas heater temperature to 400 °C, capillary temperature to 350°C, auxiliary gas flow to 12 arbitrary units, nebulizing gas flow rate to 50 arbitrary units. For quantitative analysis, a seven-point calibration curve with internal standardization was used. Tracefinder 5.1 General Quan (Thermo Fisher Scientific) was used for LC-MS data processing and metabolite quantification. Every reported metabolite was identified at level A using an authentic standard compound previously mapped to the analytical system.

### Determining the effect of pH adjustment on NAC dimerization

The effect of pH on NAC dimerization was determined at for 200 and 2 mM NAC in TSB because pH of unadjusted 200 mM solutions was below NAC’s pKa, whereas pH did not significantly differ for 2 mM solutions. 50 μL aliquots were incubated at 37°C for up to 24 hours, and freshly prepared aliquots were frozen at −80°C as a reference. After incubation, all samples were stored at −80°C. Before measurement, 10 μL sample was mixed with 80 μL methanol and 10 μL isotopically labelled internal standard, briefly vortexed and centrifuged at 10,000 rcf for 5 minutes. Supernatants were transferred to HPLC vials for measurement. For quantification, NAC standards were prepared in the range of 10 – 0.01 mg/mL in methanol with isotopically labelled internal standard to create a 7-point calibration curve. LC-MS method was run as described for bacterial samples. Skyline (V 25.1.0.237) was used for data processing, NAC and degradation products were identified using mass to charge ratios of each compound.

### Statistical analysis

Growth curves, CFU counts and NAC dimerization results were plotted using Graph Pad Prism 9.0. Before plotting of growth curves, absorbance of medium blanks was subtracted. NAC dimerization was tested with two-way ANOVA for every time point and p-values corrected with Šídák’s multiple comparisons test.

Metabolomics data was analyzed with MetaboAnalyst 6.0 (53) and Graph Pad Prism 9.0. Prior to all analysis, metabolites with >50% missing values were removed, and a 5% interquartile range filter was applied. For remaining metabolites, missing values were replaced with 1/5 of the minimum recorded value for each metabolite as missing values in our targeted LC-MS workflow stem from concentrations under the limit of detection (54).

Comparisons within one growth mode were done without normalization as bacterial counts and metabolite abundances did not differ significantly (Fig. S4). Data was Log10 transformed and pareto scaled for PCA visualization and correlation analysis. Linear correlation between NAC molarity and metabolite abundances was determined using Pearson r, significance cutoff was set at a false discovery rate (FDR) of 0.05. Abundances for intra- and extracellular metabolites were plotted from original concentrations, expressed as μM.

Median normalization was applied to compare across planktonic and biofilm growth to account for varying bacterial counts (Fig. S4) and measured metabolite abundances between both growth modes. Data was log10 transformed and pareto scaled for visualization and analysis. Planktonic and biofilm bacteria without NAC supplementation were compared to establish baseline differences, a significance threshold of FC > 1.5 and p-value < 0.1 was used. Normalized data was used for heatmap generation to compare NAC conditions across growth modes.

## Acknowledgements

This research was funded by the European Union’s Horizon 2020 research and innovation program under grant agreement No. 857287 (BBCE) and the European Union’s Recovery and Resilience Facility project Nr. 5.2.1.1.i.0/2/24/I/CFLA/003 under the grant agreement No. 1022.

We thank Fintan T Moriarty for the valuable troubleshooting and input on experimental design.

The authors acknowledge the use of Claude (Anthropic) for feedback on language and readability during the preparation of this manuscript.

## Author contribution

T.S.: Conceptualization, Methodology, Investigation, Formal analysis, Visualization, Writing – original draft, Funding acquisition. C.S.: Conceptualization, Methodology, Investigation, Writing – review & editing. V.N.: Writing – original draft, Formal Analysis. A.B: Methodology. K.K.: Supervision, Writing – review & editing, Funding acquisition. All authors have read and approved the final version of the manuscript.

## Data availability

The data sets supporting the conclusion of this article are included in the article and its supplementary files. The metabolomics dataset is publicly available at the MetabolomicsWorkbench (55) database under the project DOI http://dx.doi.org/10.21228/M8FR9X.

## Conflict of interest

The authors declare no conflict of interest.

## Supplementary material

**Fig. S1.**
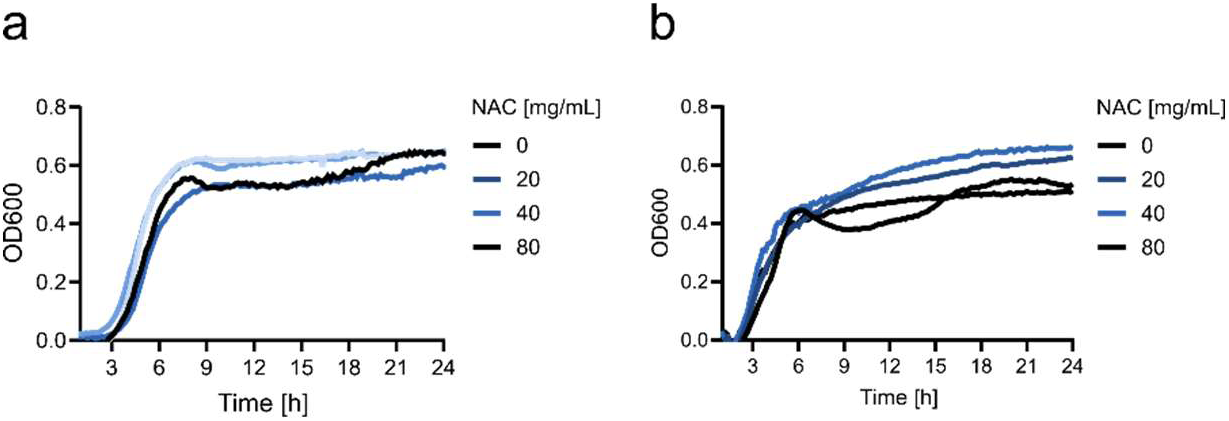
NAC at neutral pH does not reduce bacterial growth at high concentrations. Growth curves obtained by OD_600_ measurement over 24 hours in TSB supplemented with NAC close to its solubility maximum (80 mg/mL ≈ 490 mM) and further dilutions (40 mg/mL ≈ 245 mM, 20 mg/mL ≈ 123 mM) for (a) Mu12 and (b) USA300. Data represent mean of three replicates.

**Fig. S2.**
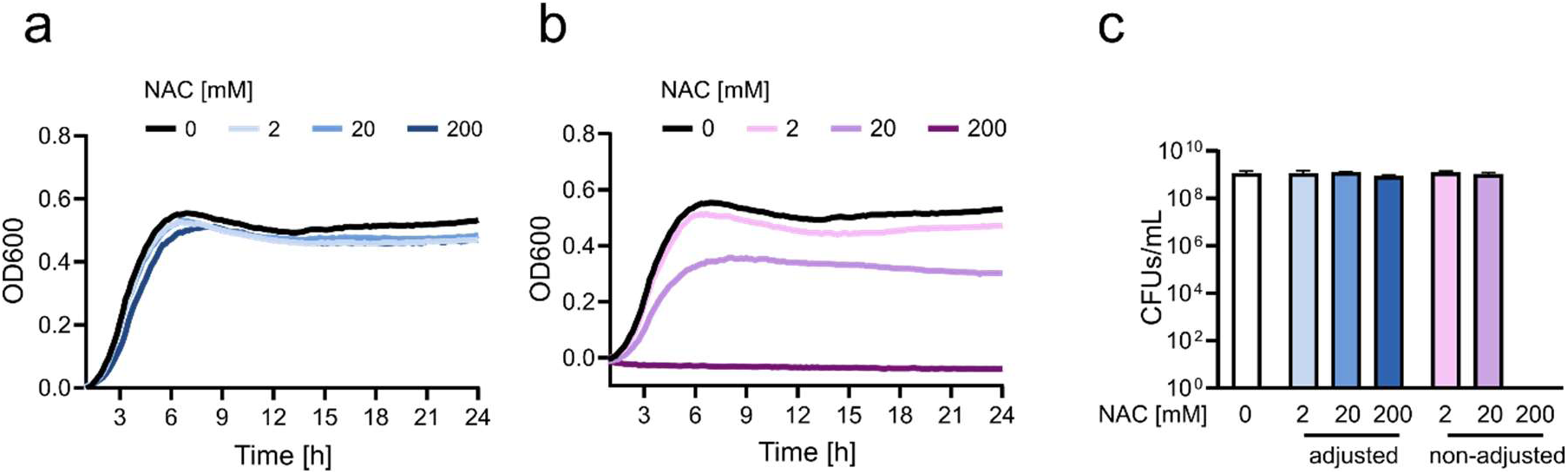
Antibacterial activity of NAC against *S. aureus* USA300 is pH-dependent, consistent with the clinical isolate Mu12. Growth curves obtained by OD_600_ measurements over 24 hours in TSB supplemented with 0, 2, 20, or 200 mM of NAC (a) with prior adjustment to pH 7 or (b) without pH adjustment. Curves represent the mean of three independent experiments. (c) Bacterial re-growth assessed by over-night CFU count of bacteria following transfer to fresh TSB after 24-hour exposure to 0, 2, 20, or 200 mM NAC with or without pH-adjustment to 7. Data represent mean ± standard deviation (SD) of three technical replicates.

**Fig. S3.**
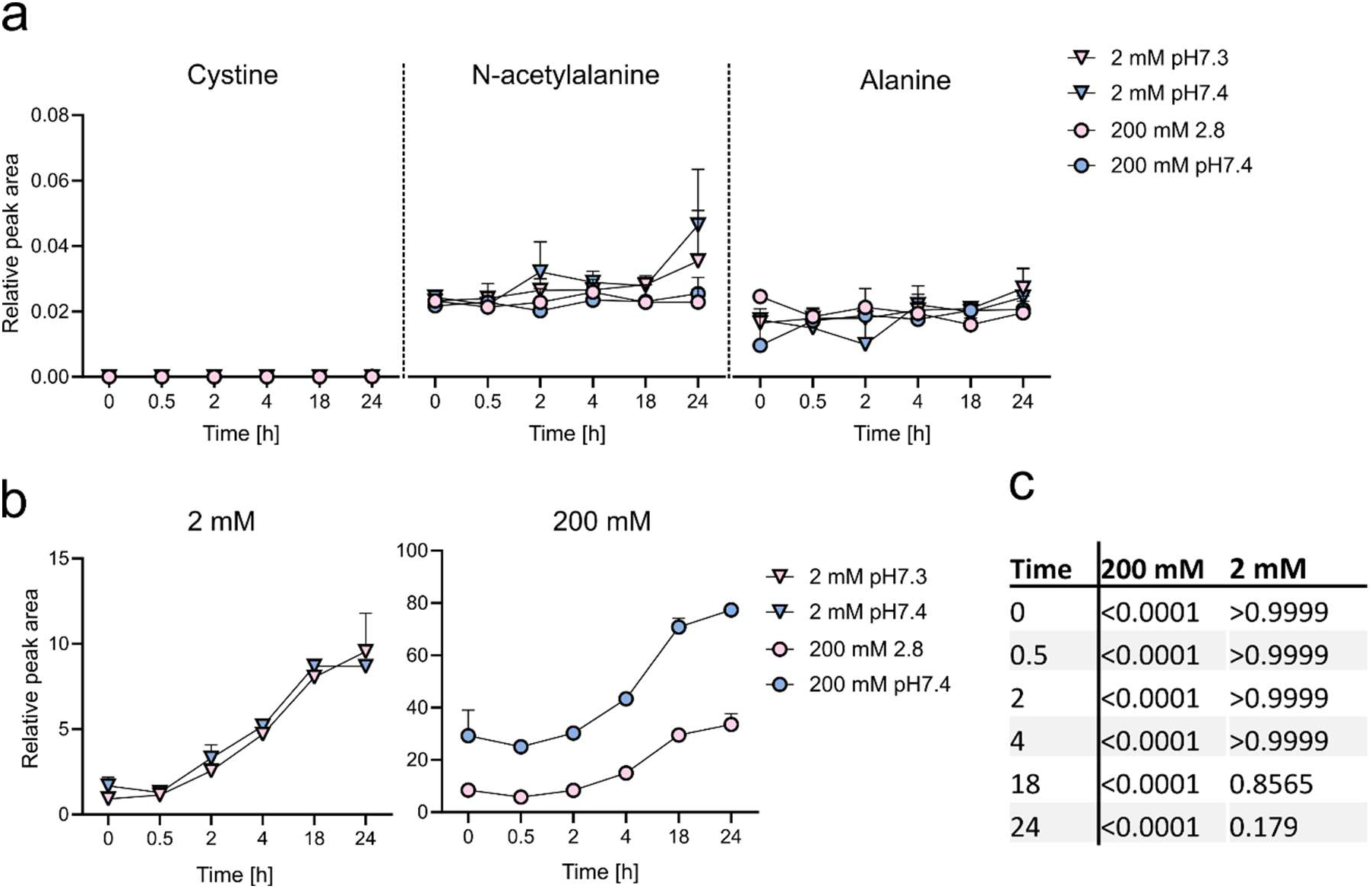
Conditions do not induce NAC decomposition but impact oxidation Sterile NAC solutions with and without pH-adjustment were incubated in TSB at 37°C for up to 24 hours to quantify NAC and its degradation products. Relative peak areas of (a) the possible degradation products cystine (deacetylation), N-acetylalanine (desulfurization), and alanine (deacetylation and desulfurization); (b) the NAC dimerization product diNAC. Relative peak areas were calculated by normalizing maximum peak areas with the internal phenylalanine standard. (c) adjusted p-values for comparison of dimer to monomer (diNAC/NAC) ratios between solutions with original and adjusted pH at every time point. p-value calculated using two-way ANOVA, corrected with Šidák’s test. Data represents mean ± SD of 3 replicates.

**Fig. S4.**
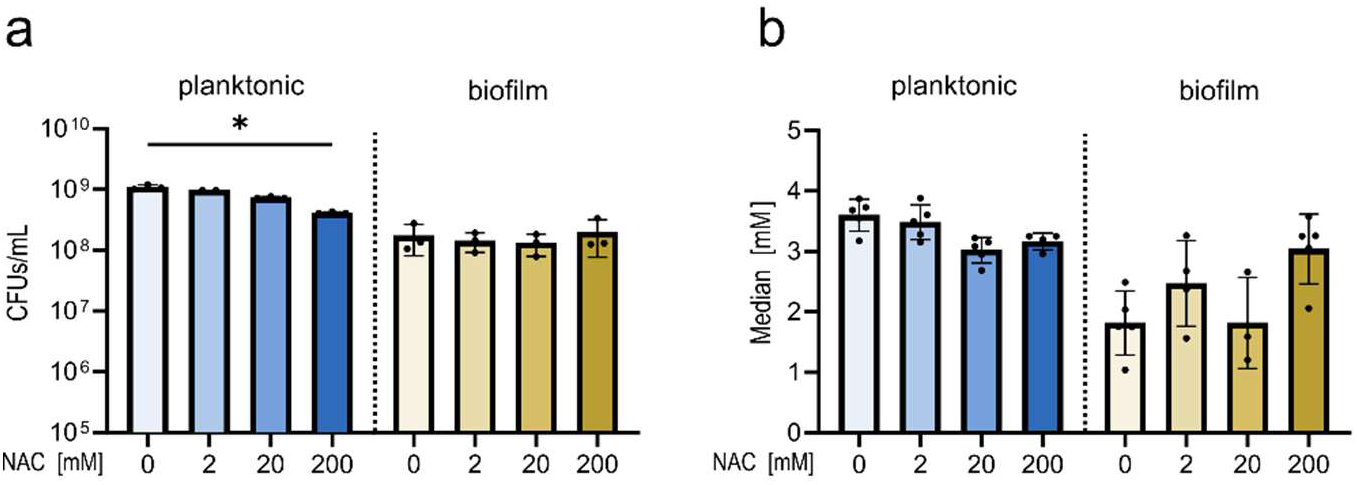
CFUs and metabolite abundances do not differ with NAC addition within growth modes. (a) CFUs/mL after growth in 0-200 mM NAC for 16 hour planktonic and 3-day biofilm cultures. 3 replicates per group used for quantification. (b) Sample medians of metabolomic measurements in 5 replicates. Significance was determined within growth modes using Kruskal-Wallis test, adjusted p-value < 0.05 used as significance cutoff.

**Table S.1.** Complete list of significant correlations and statistics of intracellular metabolite abundance and NAC concentration under planktonic growth. Significance threshold of FDR < 0.05 used.

| Metabolite` | correlation | FDR |
| --- | --- | --- |
| Ornithine | -0.97966 | 1.69E-11 |
| alpha-Ketoglutaric acid | -0.96727 | 3.09E-10 |
| Glutamic Acid | -0.94097 | 2.98E-08 |
| Cystathionine | -0.93985 | 2.98E-08 |
| Methionine | -0.92245 | 2.03E-07 |
| AMP | -0.91059 | 4.89E-07 |
| Ribose | -0.90616 | 6.46E-07 |
| Taurine | -0.89257 | 1.59E-06 |
| Acetylcarnitine | -0.88957 | 1.81E-06 |
| Aminoisobutyric Acid | -0.88857 | 1.81E-06 |
| UMP | -0.88511 | 2.15E-06 |
| Pyruvate | -0.78307 | 0.000242 |
| Erythrose 4-phosphate | -0.78183 | 0.000242 |
| Aspartic Acid | -0.77531 | 0.000275 |
| Sarcosine | -0.77474 | 0.000275 |
| Proline | -0.77411 | 0.000275 |
| Glutamine | -0.75107 | 0.000552 |
| Creatinine | -0.68049 | 0.00273 |
| Tyrosine | -0.64132 | 0.005712 |
| 4-Hydroxyproline | -0.61427 | 0.009252 |
| Aminoadipic Acid | -0.59076 | 0.013539 |
| Amino-n-butyric Acid | -0.54721 | 0.024741 |
| Adenosine | 0.50638 | 0.042434 |
| Putrescine | 0.57297 | 0.017142 |
| Glucose 6-phosphate | 0.58613 | 0.014229 |
| Carnosine | 0.64344 | 0.005643 |
| Threonine | 0.65968 | 0.004172 |
| Glucose | 0.69777 | 0.00188 |
| Serine | 0.70152 | 0.001774 |
| Asparagine | 0.71046 | 0.001468 |
| Histamine | 0.71848 | 0.001236 |
| Methylhistidine | 0.7465 | 0.000585 |
| Guanosine | 0.74928 | 0.00056 |
| Isoleucine | 0.80478 | 0.000114 |
| Alanine | 0.81327 | 8.51E-05 |
| Histidine | 0.86116 | 8.66E-06 |
| Leucine | 0.87673 | 3.55E-06 |
| beta-Alanine | 0.89752 | 1.19E-06 |
| Cystine | 0.9137 | 4.18E-07 |
| Arginine | 0.97671 | 2.66E-11 |

**Table S.2.**
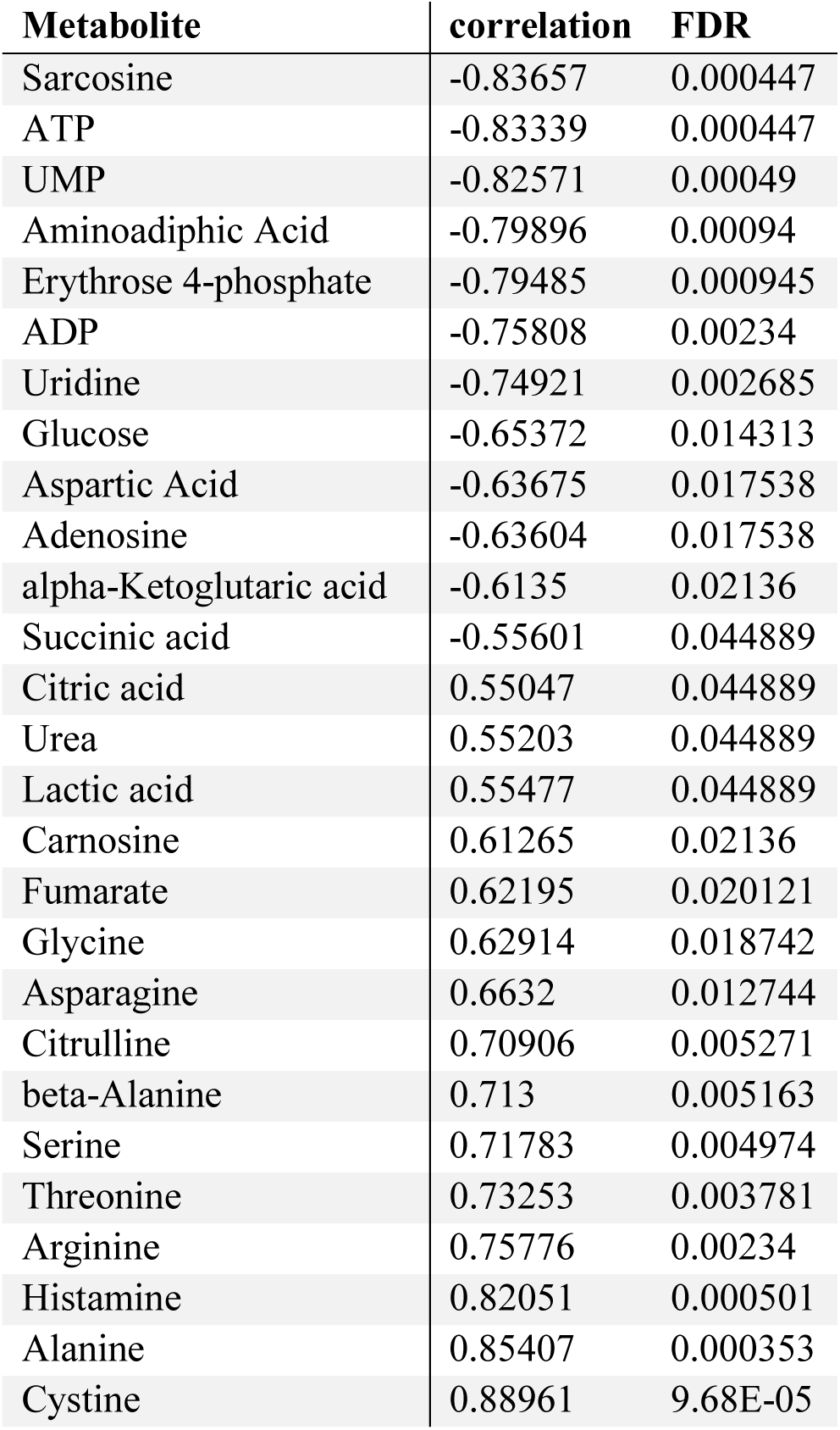
Complete list of significant correlations and statistics of intracellular metabolite abundance and NAC concentration under biofilm growth. Significance threshold of FDR < 0.05 used.

| Metabolite | correlation | FDR |
| --- | --- | --- |
| Sarcosine | -0.83657 | 0.000447 |
| ATP | -0.83339 | 0.000447 |
| UMP | -0.82571 | 0.00049 |
| Aminoadipic Acid | -0.79896 | 0.00094 |
| Erythrose 4-phosphate | -0.79485 | 0.000945 |
| ADP | -0.75808 | 0.00234 |
| Uridine | -0.74921 | 0.002685 |
| Glucose | -0.65372 | 0.014313 |
| Aspartic Acid | -0.63675 | 0.017538 |
| Adenosine | -0.63604 | 0.017538 |
| alpha-Ketoglutaric acid | -0.6135 | 0.02136 |
| Succinic acid | -0.55601 | 0.044889 |
| Citric acid | 0.55047 | 0.044889 |
| Urea | 0.55203 | 0.044889 |
| Lactic acid | 0.55477 | 0.044889 |
| Carnosine | 0.61265 | 0.02136 |
| Fumarate | 0.62195 | 0.020121 |
| Glycine | 0.62914 | 0.018742 |
| Asparagine | 0.6632 | 0.012744 |
| Citrulline | 0.70906 | 0.005271 |
| beta-Alanine | 0.713 | 0.005163 |
| Serine | 0.71783 | 0.004974 |
| Threonine | 0.73253 | 0.003781 |
| Arginine | 0.75776 | 0.00234 |
| Histamine | 0.82051 | 0.000501 |
| Alanine | 0.85407 | 0.000353 |
| Cystine | 0.88961 | 9.68E-05 |

**Table S.3.**
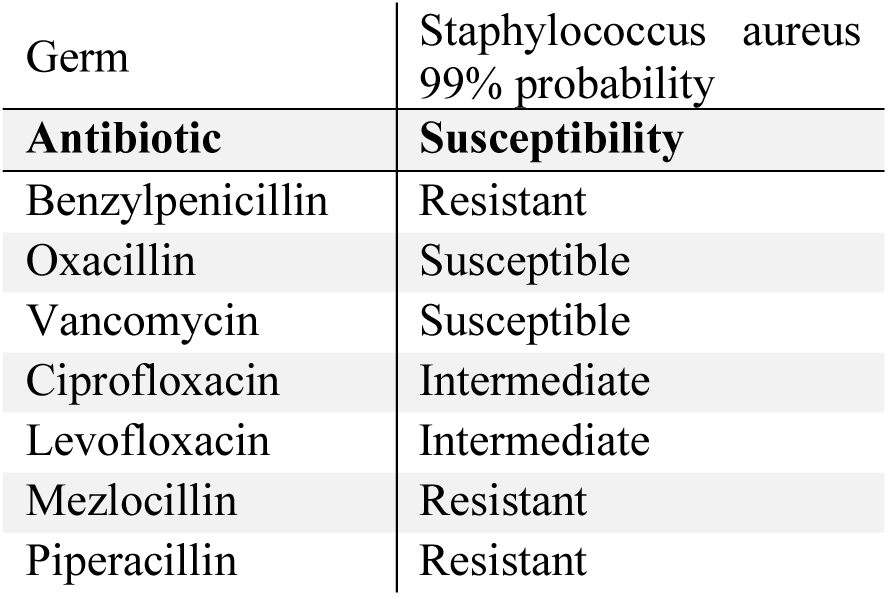
Antibiogram of the clinical *S. aureus* isolate Mu12.

